# Alternation emerges as a multi-modal strategy for turbulent odor navigation

**DOI:** 10.1101/2021.12.14.472675

**Authors:** Nicola Rigolli, Gautam Reddy, Agnese Seminara, Massimo Vergassola

**Affiliations:** Department of Physics and INFN Genova, University of Genova, Italy; Institut de Physique de Nice, Université Côte d’Azur, Centre National de la Recherche Scientifique, Nice, France; NSF-Simons Center for Mathematical & Statistical Analysis of Biology, Harvard University, Cambridge, MA, USA; Physics & Informatics Laboratories, NTT Research, Inc., Sunnyvale, CA, USA; Center for Brain Science, Harvard University, Cambridge, MA, USA; Department of Civil, Chemical and Mechanical Engineering, University of Genova, Italy; Laboratoire de physique de l’École Normale Supérieure, CNRS, PSL Research University, Sorbonne Université, Paris, France

## Abstract

Foraging mammals exhibit a familiar yet poorly characterized phenomenon, “alternation”, a momentary pause to sniff in the air often preceded by the animal rearing on its hind legs or raising its head. Intriguingly, rodents executing an olfactory search task spontaneously exhibit alternation in the presence of airflow, suggesting that alternation may serve an important role during turbulent plume-tracking. To test this hypothesis, we combine fully-resolved numerical simulations of turbulent odor transport and Bellman optimization methods for decision-making under partial observability. We show that an agent trained to minimize search time in a realistic odor plume exhibits extensive alternation together with the characteristic cast-and-surge behavior commonly observed in flying insects. Alternation is tightly linked with casting and occurs more frequently when the agent is far downwind of the source, where the likelihood of detecting airborne cues is higher relative to cues close to the ground. Casting and alternation emerge as complementary tools for effective exploration when cues are sparse. We develop a model based on marginal value theory to capture the interplay between casting, surging and alternation. More generally, we show how multiple sensorimotor modalities can be fruitfully integrated during complex goal-directed behavior.

---

The behavior of dogs alternating between sniffing in the air and close to the ground while tracking an odor scent is familiar to any cynophilist [1–4]. A similar behavior is well documented for rodents, where the slowdown associated with sniffing in the air can lead to stopping and rearing of the animal on its hind legs [5, 6]. This “alternation” between the two sensorimotor modalities strongly suggests that both airborne and ground odor cues may be exploited by animals and integrated into a multi-modal navigation strategy.

Despite the behavior’s familiarity, the reasons underlying the alternation between airborne and ground odor cues as well as the rationale of their integration are largely unknown [7]. Rodents may rear on their hind legs for a variety of reasons, generally associated with novelty detection, information gathering, anxiety and fear, as reviewed by [8]. In the laboratory odor-guided search developed by [6], mice tend to pause and rear more often in the early stages of the task. This empirical observation is consistent with rearing in response to novelty and the hypothesis that raising their head may provide the animals additional olfactory information [8]. On the physical side, it is expected that ground and airborne odor signals convey complementary information even if both signals are generated by a single source of odors. Indeed, airborne odors are valuable as distal cues because they are transported rapidly over long distances by flows that are often turbulent. The downside of airborne cues is that turbulence breaks odor plumes in discrete pockets, which can only be detected sparsely [9–12]. Furthermore, since local gradients are randomized in relation to the source direction at the timescales of olfactory searches, gradient-ascent navigation strategies are not possible [13]. Conversely, odor cues close to the ground are smoother and more continuous than odors in the air [14, 15]. The physical reason is that viscous effects make fluids slow down while flowing close to the ground at rest. As a result, boundary layers are created and the structure of the flow depends on the height from the ground [16]. In short, airborne cues are more sparse and difficult to exploit for navigation than ground signals, yet they are faster and cover longer ranges. It is therefore likely that the relative value of sniffing closer *vs* farther from the ground depends on the position of the searcher relative to the source via the statistics of odor detections that the searcher experiences. The corresponding decision of the most appropriate sensorimotor modality in response to a given history of detections is then expected to play a major role in determining an effective navigational strategy.

Here, we propose a normative theory to rationalize alternation behavior and the integration of airborne and ground-based olfactory modalities. First, we create a well-controlled setup using fully-resolved numerical simulations of the odor concentration field generated by an odor source in a channel flow. Simulations produce realistic odor plumes over distances of several meters to the source. Second, we ask what is the optimal strategy to reach the olfactory source (target) as identified by machine learning methods. Specifically, we formalize the olfactory search problem as a Partially Observable Markov Decision Process (POMDP) and use state-of-the-art methods to solve the corresponding Bellman optimization problem. The agent performing the olfactory search is given the choice between the actions of freely moving while sniffing on the ground or stopping and sniffing in the air. Solving the POMDP yields a policy of actions taken in response to a history of odor stimuli, which is encoded into a set of probabilistic beliefs about the location of the source. While the searcher could *a priori* reach the target using ground cues only, we demonstrate that learned strategies generically feature alternation between airborne and ground odor cues. Alternation is more frequent far downwind of the source and is associated with casting. The emergence of this non-trivial behavior is rationalized as the need to gather information under strong uncertainty from distal airborne cues, which leads to better long-term reward compared to local exploration for the source or proximal ground cues.

## MODEL

Consider a food source located outdoors which exudes odor at a constant rate. The odor is steadily carried by the wind and dispersed due to turbulent fluctuations. In the atmosphere, turbulent transport of odors dominates molecular diffusion and determines the statistics of the odor signal. A plume-tracking agent which enters the area downwind has to navigate its way upwind towards the source by sniffing the ground or pausing to sniff in the air for odor.

The statistics of odors on the ground is profoundly different from the statistics of odors in the air (see representative time courses in Figure 1). In the situation represented in Figure 1, the divide between air and ground is dictated by the fluid dynamics in the boundary layer close to the ground. In our direct numerical simulations of odor transport, the air travels in a channel from left and hits an obstacle at 25 cm/s, which generates turbulence. The simulations are designed to resolve the dynamics at all relevant scales, from few *mm* to several *m*, which demands massive computational resources (see Materials and Methods for details of the numerical scheme). Odor is released from a spherical source of size 4 *cm* located 56 *cm* above the ground. At the height of the source, odor is efficiently carried several meters downwind within pockets of odor-laden air which remain relatively concentrated, but are distorted and broken by turbulence. Thus odor in the air is intense but intermittent, i.e., it varies abruptly in time. Conversely, odor near the ground is smoother but also less intense (see Figure 1). It is smoother because the air in contact with the ground is still, which creates a nearly stagnant boundary layer where the disruptive effect of turbulence is tamed ; it is less intense because odorant molecules generally bind to surfaces, which act as odor sinks (see comprehensive discussions in the context of the design of olfactory tasks [17]).

**FIG. 1.**
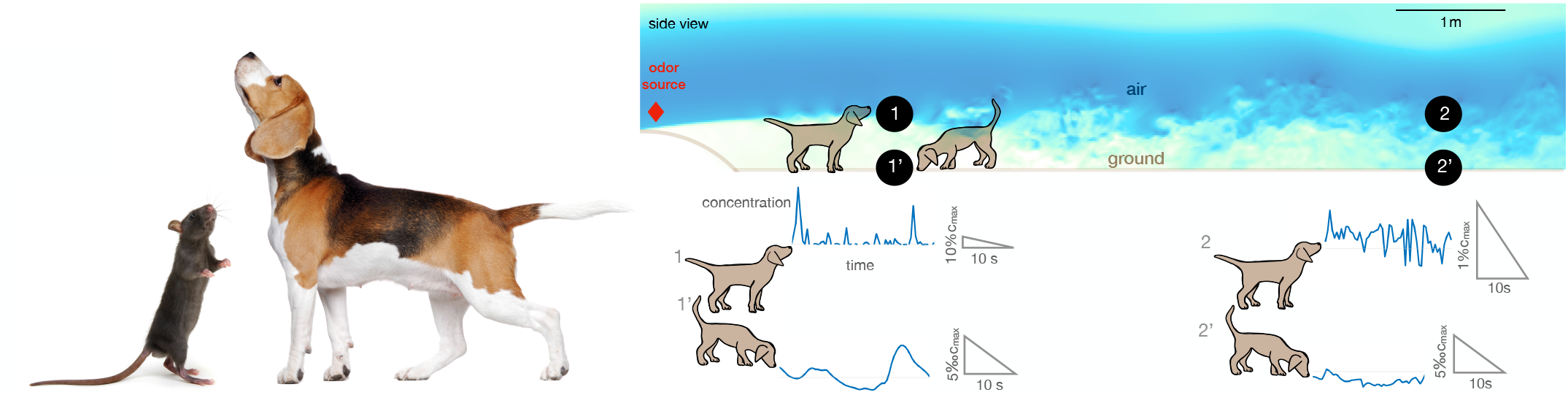
Alternation between different olfactory modalities is widespread in animal behavior. Left: a rodent rearing on hind legs and smelling with its nose high up in the air; a dog performing a similar behavior. Credit: irin-k/Shutterstock.com and Kasefoto/Shutterstock.com. Right: Side view of the direct numerical simulation of odor transport. Shades of blue represent the intensity of velocity fluctuations and are used to visualize the boundary layer near the bottom, where the velocity is reduced by the no-slip condition at the ground. Representative time courses of intense intermittent odor cues in air (sampled at 53 cm from the ground, locations marked with 1 and 2) *vs* smoother and dimmer cues near ground (sampled at 5 mm from from the ground, locations marked with 1’ and 2’). Different animals sniff at different heights, which alters details of the plumes but does not affect the general conclusions. Data obtained from direct numerical simulations of odor transport as described in the text, see Methods for details.

Our simulations specifically consider the limiting case of total adsorption of odor molecules. Qualitatively similar results are expected for models intermediate between total adsorption and total reflection, where particles have a finite likelihood of being adsorbed or re-emitted in the bulk. Total (or partial) depletion of odors at the ground surface implies that an agent with a finite detection threshold can only sense ground odors near the source. Conversely, the agent is able to detect odor in air across a more extended area (compare top views in the air *vs* the ground in Figure 2(a)-(b)), that is agents can sense larger plumes in the air than on the ground (Figure 2(c)).

**FIG. 2.**
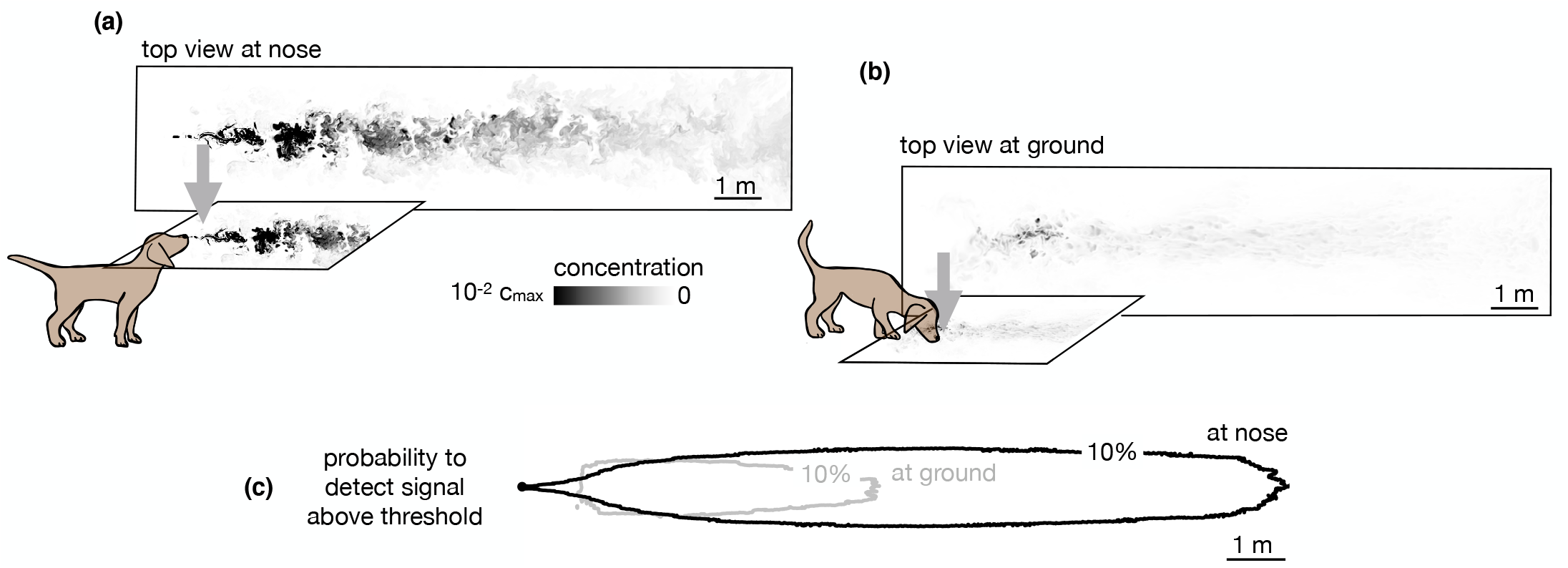
Snapshots of odor plume obtained from direct numerical simulations of the Navier-Stokes equations in three spatial dimensions. Top view of the odor plume **(a)** at nose height and **(b)** at ground. **(c)** 10% isoline of the probability to detect the odor *r*(*x, y*) (defined as the probability that odor is above a fixed threshold of 0.14% with respect to the maximum concentration at the source) at the ground (grey) and at the nose height (black).

The statistics of odor encounters detected along the search path of a plume-tracking agent provides useful information about the location of the source, which guides subsequent navigation. We consider an agent moving along a path *r*_1_, *r*_2_, …, *r*_*t*_ while measuring the odor signal *o*_1_, *o*_2_, …, *o*_*t*_. The agent’s present knowledge is fully summarized by the posterior distribution of the agent’s location relative to the source, ***b***_*t*_, also called the belief vector. The agent computes Bayesian updates of the belief ***b***_*t*_ using a model (the likelihood of odor detections) and the current observation *o*_*t*_. At each time step, *t*_*s*_, the agent decides among six alternatives: staying at the same location or moving to one of the four neighboring locations while sniffing the ground, or staying at the same location and sniffing in the air. Choices are dictated by the long-term reward that the agent expects to receive, as discussed in the next paragraph.

We pose the agent’s task in the framework of optimal decision-making under uncertainty. The agent’s actions are driven by a unit reward received when it successfully finds the source. Rewards are discounted at a rate *λ*, i.e., the expected long-term reward is ⟨*e*^−*λT*^ ⟩_*T*_. Here, *T* is the time taken to find the source and the expectation is over the prior knowledge available to the agent, its navigational strategy and the statistics of odor encounters. The expected long-term reward or *value, V* (***b***_*t*_), given the current state of knowledge, ***b***_*t*_ can be calculated using the Bellman equation, a dynamic programming equation which takes into account all possible future trajectories of an optimal agent. Specifically, we obtain the Bellman equation (see Methods for details)

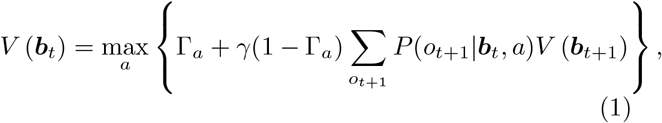

where Γ_*a*_ is the probability of finding the source immediately after taking action 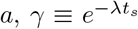 and the probability *P* (*o*_*t*+1_|***b***_*t*_, *a*) of observing *o*_*t*+1_ is determined by the physical environment and the signal detection threshold of the agent. Intuitively, the terms in the argument of the max function in (1) represent the value of finding the source, detecting the odor signal or not detecting the odor signal, each event being weighted by its probability. The optimal action is the one that maximizes the value, i.e., the parenthesis on the right-hand side of (1). For simplicity, we discretize observations into detections (odor signal above a fixed threshold) and non-detections, which implies that the behavior depends solely on the probability per unit time of detecting the odor on the ground or in the air (Figure 2c). Thus, the agent uses a (partially inaccurate) Poissonian detection model, whose maps of the average detection rate do match the simulated odor plumes but they lack the appropriate spatiotemporal correlations. See Results for more details and the corresponding performance.

The decision-making dynamics form a partially-observable Markov decision process (see Methods for a brief introduction to POMDPs). To solve (1), we use approximate methods, which exploit a piecewise linear representation of the value function: *V* (***b***_*t*_) ≈ max_*i*_{***α***_*i*_.***b***_*t*_}. The ***α***_*i*_’s are a set of hyperplanes, which are found using (1) along exploratory search paths and yield increasingly accurate approximations of the value function with training. For each trial run, we begin with a uniform prior distribution and simulate the POMDP until the agent finds the source. If the agent does not find the source within 1000 steps, we interrupt the simulation. The time step, *t*_*s*_, and the distance traveled at each step are set such that the agent sniffs three times per second and at every step it moves 12 cm.

## RESULTS

### The agent navigates by alternating between sniffing the ground and air

An agent initially downwind of the odor source learns to navigate the odor plume to maximize the discounted reward described previously, that is, to minimize the search time. The upshot of the learning phase is that the final search policy alternates between sniffing on the ground and the air (Figure 3). The average time taken to reach the source reduces considerably with the training time, indicating the emergence of an effective navigational strategy (Figure S1).

**FIG. 3.**
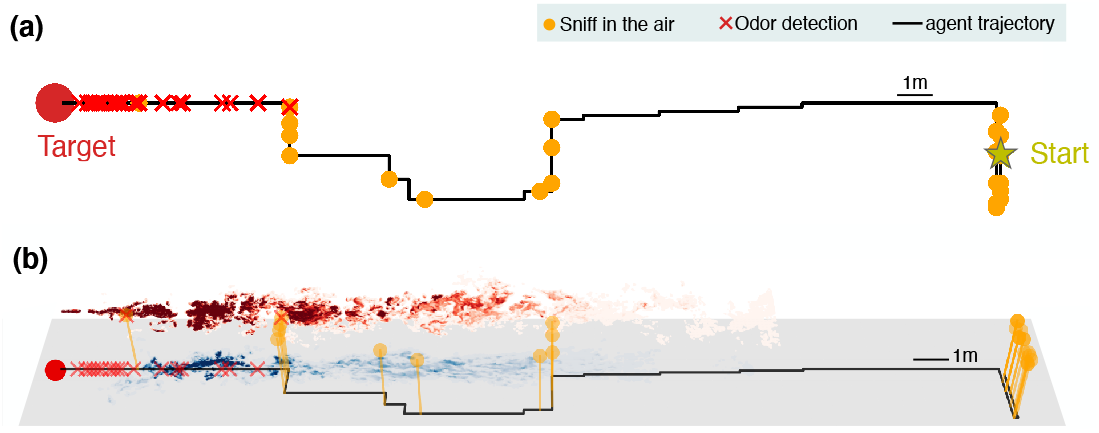
Representative trajectories undertaken by an agent learning how to reach the source of a turbulent odor cue. **(a)** Top view of a representative trajectory at the end of training. **(b)** Three dimensional view of sample trajectory from panel (a), superimposed to two snapshots of odor plumes near ground (shades of blue) and in the air (shades of red). Trajectories are obtained by training a POMDP, where the agent computes Bayesian updates of the belief using observations (odor detection or no detection) and their likelihood (detection rates from simulations of odor transport). Agents trained with this idealized model of odor plumes successfully track targets when tested in realistic conditions (see Movie S1).

The trajectories learnt by the agent display a variety of behaviors reminiscent of those exhibited by animals, which include wide crosswind casts interleaved with up-wind surges. Notably, the agent exhibits a recurring motif which cycles between moving to a new location and pausing to sniff in the air. The alternating behavior emerges directly as a consequence of the statistics of the physical environment in spite of pausing to sniff in the air, which leads to the cost of a stronger discount in the reward.

When, where and why does the agent sniff in the air? Trajectories shown in Figure 3 exhibit extensive alternation at the beginning of the search when the agent is far downwind compared to when it is close to the source. A quantitative analysis across training and test realizations confirms that the agent’s rate of sniffing in the air is significantly higher farther away from the source (Figure 4(a)). This observation is rationalized by the greater probability of detecting an odor signal in the air at distant locations (Figure 2(c)) despite the increased intermittency in the airborne signal (Figure 1). In spite of the added cost entailed by slowing down locomotion, sniffing in the air ultimately speeds up the localisation of the source (Figure 4(b)). This behavior is maintained across different training realizations and when the discount factor, *γ*, is reduced so that the delay incurs a greater cost. In sum, alternation emerges as a robust, functional aspect of an effective long-term strategy of olfactory search (see also Figure S2).

**FIG. 4.**
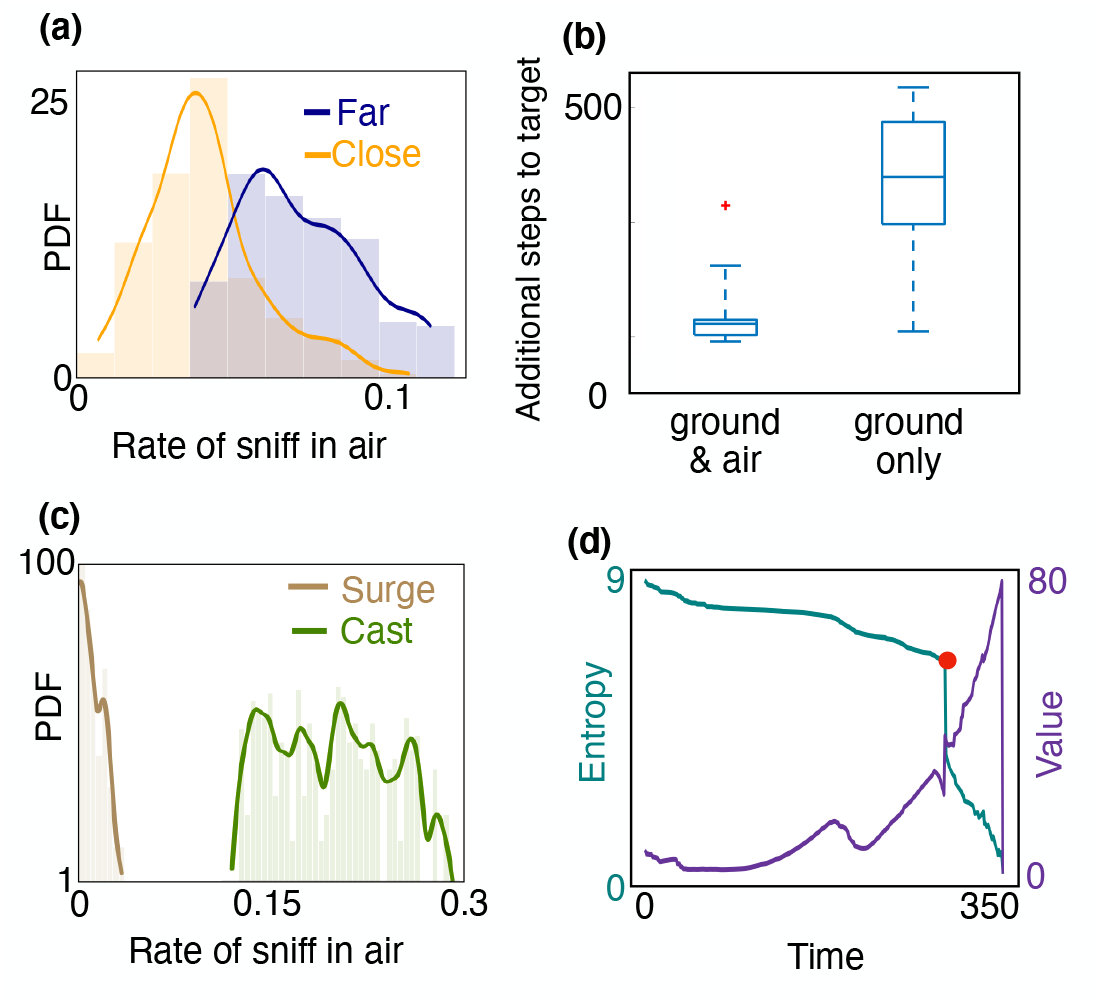
Empirical characterisation of the alternation between olfactory sensory modalities. (**a**) The agent sniffs more often in the air when it is far from the source, i.e., outside of the airborne plume. The rate of sniffing in the air is the fraction of times the agent decides to sniff in the air rather than move and sniff on the ground. The fraction is computed over the entire trajectory in the conditions identified in the different panels. Statistics is collected over different realizations of the training process and many trajectories, with different starting positions (see Materials and Methods for details). **(b)** The number of steps needed to reach the target minus the number of steps needed to travel from the starting position to the source in a straight line. The horizontal line marks the median, boxes mark 25^*th*^ and 75^*th*^ percentiles; red dot: outlier (value exceeds 75^*th*^ percentile + 1.5 × interquantile range). Dashed lines mark 10^*th*^ and 90^*th*^ percentile. For reference, a straight line from the center of the belief to the source is 240 steps. Agents that are given the possibility to pause and sniff in the air are able to reach the target sooner than agents that can only sniff on the ground. **(c)** Agents sniff in the air once every 5 steps on average when they cast, whereas they only sniff in the air once every 60 steps while surging upwind. We consider the agent to be surging if it moves *k* consecutive steps upwind and casting if it moves *k* consecutive steps crosswind or sniffs in the air. We use *k* = 3, results shown hereafter do not depend strongly on this choice. **(d)** Entropy (cyan) and value (purple) of the belief *vs* time, along the course of one trajectory. The red dot indicates a detection, which provides considerable information about source location and thus makes entropy plummet and value increase.

A striking feature of the trajectories in Figure 3 is the strong correlation between casting and sniffing the air before the first detection is made. To quantify this effect, we categorize the agent’s behavior into casts (persistent crosswind movements) and surges (upwind movement), and measure the rate of sniffing the air for both of these behaviors. We find that the rate of sniffing in the air is typically an order of magnitude greater during casts as compared to surges (Figure 4(c)), indicating that alternation is tightly linked to the switch between casting and surging. Casting has been classically interpreted as a strategy for efficient exploration in an intermittent environment. The coupling between casting and alternation observed here suggests that sniffing in the air is an alternative mode of exploration which aids and complements casting when searching for a sparse cue. Exploration dominates the first part of the search until the the first detections which substantially reduce uncertainty (see entropy of the posterior distribution in Figure 4(d)).

Overall, we are led to the following picture of the search dynamics. At the beginning of the search, the agent has a broad prior that is much larger than the odor plume of size ∼ *x*_thr_ × *y*_thr_, where *x*_thr_ is the plume length and *y*_thr_ is the plume width when sniffing in the air. The agent then has to identify and home into the *x*_thr_ × *y*_thr_ region that contains the odor plume. The bottleneck in this phase is the scarcity of odor detections, which require an efficient exploration strategy. Once the odor plume is detected, the agent knows it is near the source and the search is driven by surface-borne odor cues, while the frequency of sniffing in the air is significantly reduced. In short, our simulations show that the behavior can be split into two distinct phases: 1) an initial exploration phase accompanied by extensive casting and alternation, where the agent attempts to localize the plume, and 2) odor-guided behavior in a regime relatively rich in cues, which enable the agent to precisely locate the source within the plume.

We conclude this Section noting that the above remarks are expected to hold more generally than in the specific setup of our simulations. Movie S1 shows that the same behaviors are displayed by agents navigating a realistic plume despite their learning in a (partially inaccurate) Poissonian model of odor detections. This finding indicates the robustness of the learning scheme to inaccuracies in the model of the environment, which are inevitably present in any realistic situation. More specifically, the static information provided by the average detection rate map is found to be sufficient for navigation and alternation. While more information on dynamical spatiotemporal correlations may help further improve performance, the fundamental requirement for alternation is the presence of wider detection rate maps in the air than on the ground. Thus, as long as this feature is preserved, we expect agents to display alternating behaviors and the two phases mentioned above. In particular, these properties should hold also when different models of odor transport are employed by the searcher and/or surface adsorption chemistry is more involved than pure adsorption.

### Intuition

We now proceed to understand the transition between the two distinct phases of search that were identified previously. While a detailed analysis of each individual decision is challenging, we can gain intuition by decomposing the agent’s overall behavior into segments. Each segment is then rationalized by examining how the agent explores the locations where it believes it can find an odor signal or the source. For this purpose, let us examine how the agent’s belief of its location relative to the source evolves as the search proceeds.

In the representative example depicted in Figure 5, the agent begins with a uniform prior belief, much larger than the plume, as shown in the top row of Figure 5. The agent makes its first action by sniffing in the air and does not detect an odor signal. Since odor is not detected, the likelihood that the agent is immediately downwind of the source is reduced, which leads to a posterior belief updated via Bayes’ rule (second row, Figure 5). The agent proceeds by casting crosswind in a loop while occasionally pausing to sniff in the air (third row, Figure 5), after which it executes an upwind surge (fourth row, Figure 5). The decision to surge at that specific moment can be understood from examining the belief immediately before the surge: because the agent did not detect any odor over the entire cast-and-sniff sequence, the likelihood that the agent is located near the source, i.e., within the plume, is extremely low (third row, Figure 5). At this point, it is more valuable to surge upwind rather than continuing to explore the same area. By surging forward, the agent is now more likely to encounter the plume, which enables it to effectively explore the remaining part of the belief. The key to the above argument is that the agent lacks knowledge of its position relative to the source, and it acts so as to narrow down its belief. Indeed, over the course of a search, entropy of the belief steadily declines, while its value increases (Figure 4(d)).

**FIG. 5.**
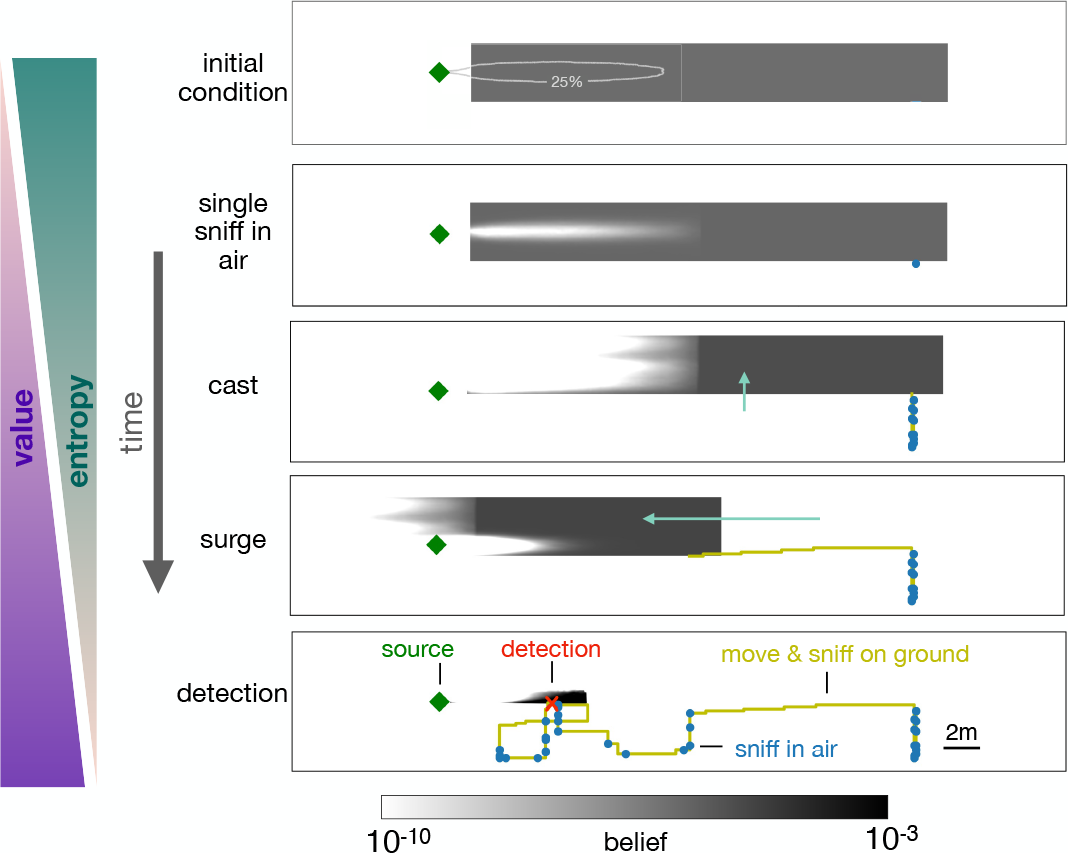
Progression of the belief the agent has about its own position relative to the source. From top, first panel: before starting the search the agent has a flat belief about its own position, much broader than the plume in air represented by the 25% isoline of the probability of detection. Second: belief after a single sniff in the air and no detection. The white region corresponds to the extent of the plume in air and indicates that because the agent did not detect the odor, it now believes it is *not* within the plume right downstream of the source. Third: As the agent casts, its belief about its own position translates sideways with it; additionally, at each sniff in the air with no detection, the belief gets depleted right downstream of the source, as in the panel right above. As a result, the cast-and-sniff cycle sweeps away a region of the belief as wide as the cast and as long as the plume. Fourth: As the agent surges upwind, its belief about its own position translates forward with it; additionally, as it sniffs on the ground with no detection, the belief gets depleted in a small region right downstream of the source, corresponding to the extent of the plume on the ground. Fifth: after detection, the belief shrinks to a narrow region around the actual position of the agent, which leads to the final phase of the search within the plume. Green (Purple) wedges indicate that the entropy of the belief decreases (value of the belief increases) as the agent narrows down its possible positions (and approaches the source).

A repetition of the sequence of casting, alternation and surging follows as the agent steadily narrows down the belief, until it finally detects the odor (bottom panel, Figure 5). The detection shrinks the posterior to a small patch which makes entropy plummet (Figure 4(d)) and leads the agent rapidly to the source. The first detection event (identified by the red dot in Figure 4(d)) is what marks the transition between searching *for* the plume and searching *within* the plume, as discussed above.

### Searching for airborne cues

We now expand on the intuition above by introducing a simplified, quantitative model of the search. Its goal is to address the search dynamics in the initial phase before detection, when the agent searches for the plume. This is the key phase as the localization of the plume largely dominates the search time (see red dot in Figure 4(d)).

To introduce the main simplification of the model, we note that in the exploratory regime at large distances, the agent is more likely to detect odor by sniffing in the air due to the larger detection range of airborne cues. We therefore ignore odor signal on the ground and assume the agent only detects odor by sniffing in the air. This simplifies the analysis considerably as the search path is then parameterized by the discrete locations at which the agent sniffs in the air rather than the specific trajectory taken between sampling locations. The prior distribution, ***b***(*x, y*), of the agent’s location with respect to the source is assumed uniform with length *L*_*x*_ (≫ *x*_thr_) (along the downwind direction) and width *L*_*y*_ (≫ *y*_thr_), similar to the example shown in Figure 5 (top). The probability of detecting an odor signal in a sniff, *r*(*x, y*), depends on the extent of the plume via the parameters *x*_thr_ and *y*_thr_. To decouple the upwind surge and cross-wind cast, we approximate the detection probability map in Figure 2(c) as *r*(*x, y*) = *f* (*x*)*g*(*y*), where *f* (*x*) is a constant when 0 < *x* < *x*_thr_ and 0 otherwise, *g*(*y*) has a characteristic length-scale *y*_thr_.

To localize the plume, the agent has to sufficiently explore, by sniffing in the air, patches of size ∼ *x*_thr_ × *y*_thr_ within its prior. Since the prior’s width is larger than the plume width (*L*_*y*_ ≫ *y*_thr_), the agent has to cast in order to determine how far it is from the plume’s center-line. Each sniff effectively explores a patch of length ∼ *x*_thr_ immediately downwind of the source. Therefore, a bout of casting across a width *L*_*y*_ while constantly sniffing in the air explores a region of size ∼ *x*_thr_ × *L*_*y*_. There, the likelihood of containing the source is strongly depleted, which converts the initial prior into a posterior of reduced length *L*_*x*_ − *x*_thr_. Since the agent now believes to be outside of the plume, it is convenient to continue the search surging upwind by *x*_thr_, and exploring a new patch via casting. The process is repeated until the plume is detected.

The search process can therefore be split into distinct episodes where the agent cycles between sniffing while casting and surging upwind by ∼ *x*_thr_. We identify three main questions about the search, which we address in more detail below: 1) how wide should the agent cast? ; 2) how long should the agent spend casting before surging upwind? ; 3) where should the agent sniff during the casting phase? Specifically, we highlight and quantify the various trade-offs associated with the cast-sniff-surge modes of exploration.

Since the rate map is uniform in *x* and has length *x*_thr_, the agent surges exactly a distance *x*_thr_. The search process is then decomposed into *N* ∼ *L*_*x*_*/x*_thr_ distinct episodes. In each episode *n* (*n* = 1, … *N*), the agent spends time *t*_*n*_ deciding whether the source is within reach, i.e., closer than *x*_thr_. The casting duration *t*_*n*_ is to be optimized. After *t*_*n*_, since the agent has determined that the source is not yet within reach, it surges upwind and continues to the next episode *n* + 1. The process continues until the agent obtains a detection. The cumulative probability of not detecting the signal (conditional on the target being in that patch) after casting for time *t, c*(*t*), depends on the sampling strategy during casting and is discussed further below.

The expected discounted reward at the beginning of the search is *V*_1_ ≡ ⟨*e*^−*λT*^ ⟩_*T*_, where *T* is the time taken to find the odor signal. We use dynamic programming to compute and optimize *V*_1_. *V*_1_ is the sum of the expected reward if the agent finds the signal in the first patch within time *t*_1_ and the expected reward after moving to the next patch if it does not. The information gained from the observation of not detecting a signal is taken into account in the latter term through a Bayesian update of the prior. However, we show that *V*_1_ and the casting times, *t*_1_, *t*_2_, …, *t*_*N*_, can be calculated using an equivalent, simpler expression which does not require Bayesian updates (Methods). Specifically, denote *V*_*n*_ as the expected discounted reward at the beginning of the *n*th episode, i.e., before the cast and surge. *V*_1_ is calculated using the recursive equation

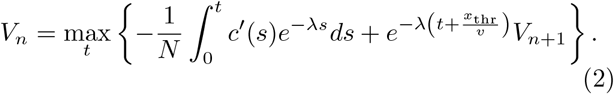

The time *t* that maximizes the parenthesis determines the optimal duration *t*_*n*_ the agent should spend casting before surging upwind. The first and second terms in the parenthesis of (2) are the expected discounted rewards if the agent detects a signal during casting (and the search ends) or if it does not detect a signal, surges a distance *x*_thr_ and continues to the next episode, respectively. The factor −*c*′ in the first term is the probability density to make a detection at time *t* conditional on the target being in the current patch, which has probability 1*/N*.

We first show that the duration *t*_*n*_ obeys a marginality condition. The agent should stop casting when the value of continuing to explore the current patch is just outweighed by the value of moving on and exploring the next patch. This intuition is quantified by optimizing for *t* in (2). Zeroing the time derivative of (2), we obtain that *t*_*n*_ is the value of *t* that satisfies the equality 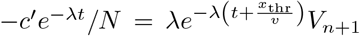. The left hand side is the rate of value acquisition upon staying in the current patch. The right hand side is the negative rate of value acquisition upon delaying departure, that is the rate of value acquisition upon anticipating departure. Thus by maximizing value we obtain that, at optimality, the added value of continuing to cast matches the added value of anticipating surge, i.e., marginality of the two actions as prescribed by marginal value theory [18]. The marginality condition leads to a relationship between the casting time and the value at the next episode

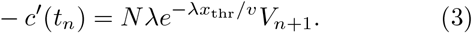

When *n* = *N*, the agent casts indefinitely, which gives 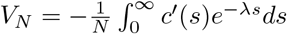 from (2). The casting time for each episode is obtained using this boundary condition, (3), and *c*(*t*), which we shall determine in the next paragraph. Note that we have ignored the possibility that at *n* = *N*, the agent turns back and moves downwind to re-explore earlier regions, which can be incorporated into this framework and leads to a different boundary condition. However, we do not take this into account since this extension only marginally affects the earlier stages of the search path and does not affect general conclusions.

We now optimize for the sampling strategy during casting, which in turn determines *c*(*t*). The casting phase can be formulated as a decision-making process of deciding where to sniff next on the crosswind axis given the marginal distribution 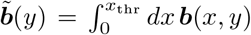. The next sniff location at a displacement Δ*y* from the current location is obtained from the dynamic programming equation similar to (2), which relates the current value to the value of moving and sampling elsewhere.

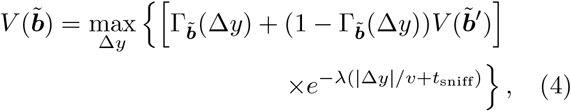

where 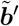 is the posterior after sampling at the new location conditional on no detection, and 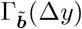 is the probability of detection. The two terms in the Bellman equation correspond to the cases when the agent detects a signal and does not detect a signal respectively, which are discounted in proportion to the time taken to travel a distance | Δ*y* | and sniff in the air. Numerically solving (4) yields a sampling strategy and the corresponding *c*(*t*). The optimized casting strategy is a zigzag (Figure 6a) which expands over time to the width of the prior. The probability of not detecting the signal decays exponentially with a rate depending on the optimization depth (Figure 6b). In the low-detection rate limit, we generically expect a constant detection rate (say *c*(*t*) = *e*^−*κt*^), consistent with the exponential decay observed in the simulations. The detection rate *κ* decreases with *t*_sniff_ (Figure 6c), which in turn translates to a decreased value (from (3)) and highlights the cost of pausing to sniff in the air. From (3), we then have

**FIG. 6.**
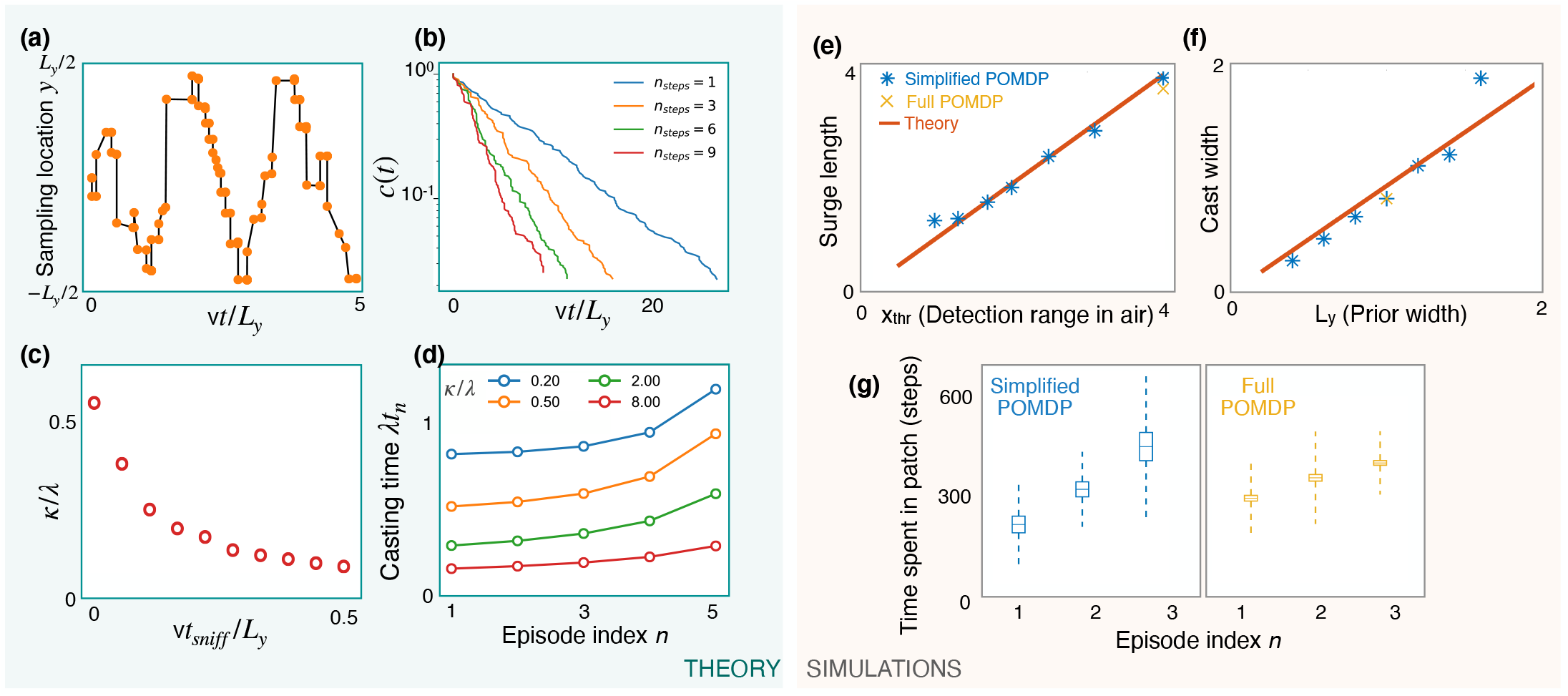
(a) Optimized sniff locations during casting (conditional on no detection) show a zigzag of increasing amplitude. At each decision, (4) is expanded and optimized w.r.t the subsequent *n*_steps_ = 10 sniff locations 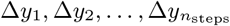 using standard black-box optimization methods. The agent then moves by Δ*y*_1_ and the procedure is repeated. (b) The probability of not detecting the signal against time, *c*(*t*), decays exponentially with detection rate, *κ*, shown here for different values of the optimization depth *n*_steps_. *κ* saturates beyond *n*_steps_ = 9. (c) *κ* monotonically decreases with the time per sniff, *t*_sniff_, reflecting the cost of pausing to sniff the air. In panels (a), (b) and (c), we use *y*_thr_*/L*_*y*_ = 1*/*20, *L*_*y*_ = 1, *λ* = 0.5, *v* = 1 and *t*_sniff_ = 0 (for (a) and (b)). (d) Casting times (in units of 1*/λ*) generally increase as the search progresses. Obtained using (5) for different values of *κ/λ* (colored lines). Here *N* = 6 and *λx*_thr_*/v* = 0.05. (e,f) The surge length and cast width from simulations of a simplified POMDP, where the agent can detect an odor signal only by sniffing in the air. Results for different prior and plume dimensions (blue stars) align with the theoretical prediction (red line) that the surge length and cast width are equal to the detection range in air, *x*_thr_, and the prior width, *L*_*y*_, respectively. Results from the full POMDP, where the agent can detect odor on the ground, are also consistent with the predictions (yellow crosses). (g) The time spent casting in each patch for the simplified and full POMDP increases as the search progresses, as predicted by the theory (panel (d)). Here we set the prior length, *L*_*x*_ = 4*x*_thr_, which corresponds to *N* = 4 patches. Boxes and dashed lines represent the standard error and the standard deviation around the mean respectively.

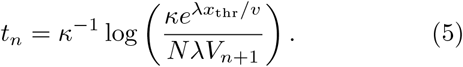

We use (2) and (5) along with the boundary condition 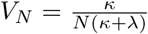 to solve for the casting times. The results show increasing casting times with episode index (Figure 6d). Intuitively, as the search progresses, the marginal cost for the agent to continue casting decreases due to its increasing confidence that it is in the right patch, driving the agent to spend more time casting before leaving the patch.

We test predictions from the theory using simulations of a simplified POMDP. Specifically, the agent is trained to find the target with an odor signal that can be detected only by sniffing in the air. The detection probability map *r*(*x, y*) is rectangular with plume detection range *x*_thr_ and width *y*_thr_. Simulations confirm that the surge length and cast width are equal to the detection range and the prior width respectively (Figure 6e,f). The time spent exploring a patch increases monotonically as the search progresses, as predicted by the theory (Figure 6d,g). Notably, we find that the trajectories from the full POMDP considered in the previous sections are also consistent with these predictions, suggesting that these aspects are generic features of foraging for a sparse odor signal during the first phase of exploration.

## DISCUSSION

Motivated by the goal of disentangling elementary components in the complexity of animal behavior, we have investigated a dynamics driven entirely by olfactory cues. In the model, an agent searches for a source of odors transported by a turbulent flow and at each step decides either to move while sniffing on the ground or pause, rear up and sniff in the air. The goal is to locate the source in the shortest possible time, which is the reward function used to identify effective policies of action using machine-learning methods. Analogously to dogs and rodents mentioned in the introduction, we obtain behavioral policies which feature alternation between the two modalities of sniffing on the ground *vs* in the air. The appeal of our approach is that we could identify the rationale for the observed alternation and its basic factors. On the one hand, movement and progression toward the source is halted during the rearing phase of sniffing in the air. On the other hand, odor sources create large turbulent plumes that reach larger distances in the air than on the ground. Therefore, sniffing in the air may have a higher chance of intersecting odor cues than on the ground. These two competing effects underlie the process of alternation and their balance determines the rate of switching between the two modalities, which depends on the distance as discussed in the next paragraph.

The effect of alternation is particularly pronounced at large distances to the source. There, due to turbulent mixing, the odor concentration drops substantially and no gradients are present [12]. In our realistic setting, where the searcher does start at large distances, the process can be qualitatively split in two phases : first, the agent needs to approach the source enough for an almost continuous odor plume to be present ; second, it needs to locate the source within the plume. The latter task, which is the regime that most laboratory experiments have considered so far [7], is much easier than the former as the rate of odor detection close to the source and within the conical plume is relatively high. Therefore, the task boils down to staying close to the center of the conical plume, where the signal is highest. Conversely, the bottleneck during the first, harder phase is the scarcity of information on the location of the source, which the agent tries to overcome by increasing its chances of odor detection. Slowing down its progression is thus the price that the agent pays in order to get oriented in the uncertain conditions typical of large distances to the source. The transition between the two search phases typically occurs after a handful of odor detections.

Note that we have focused here on the case of a stationary source, where odor statistics in the air and on the bottom layers are discriminated by the adsorption on the ground. In fact, at the onset of odor emission (and even in the absence of adsorption), plumes start out larger in the air than near the ground, simply because air travels more slowly near the ground. It follows from our results that alternation should be more frequent in the early stages of odor release in non-steady conditions. This prediction could be tested experimentally by switching on an odor source and monitoring the fraction of sniffing in the air as a function of the time elapsed since the switch and the onset of odor emission.

The machine-learning methodology that we have employed here to identify effective policies of actions belongs to the general family of Partially Observed Markov Decision Processes (POMDP) [19, 20]. This framework applies to a broad class of decision problems, where agents need to accomplish a prescribed task by a series of actions taken with partial knowledge of the environment. Specifically, the agent combines external cues and its internal model of the world to infer a belief about the state of the environment. In our setting, the agent is the searcher, cues are odor detections (or their absence), the task is to localize the source, and beliefs pertain to the location of the source of odors. While the agent proceeds along its path and gathers information via odor cues, its belief narrows down and eventually concentrates at the location of the source. Trajectories of a POMDP agent with a single sensory modality and their relation to phenomenological approaches as Infotaxis [13] were discussed in [7]. Here, we have given the agent the choice of multiple sensory modalities at each decision step, which allowed us to highlight the presence of alternation and establish its link with marginal value theory (MVT) [18].

MVT describes the behavior of an optimally foraging individual in a system with spatially separated resources. Due to the spatial separation, animals must spend time traveling between patches. Since organisms face diminishing returns, there is a moment the animal exhausts the patch and ought to leave. In MVT, the optimal departure time is determined as the time at which the marginal value of staying in a patch equals that of leaving and exploring another patch. In our setting, these patches correspond to regions of the agent’s belief which are explored using a combination of casting and sniffing in the air. MVT thus determines when to stop cast-and-sniff exploration and surge towards the next patch in the belief.

While we considered two olfactory sensorimotor modalities, our methodology and results apply more broadly to distinct sensory systems and cues. If there is no conflict in the acquisition and processing of multiple sensory cues, then it is clearly advantageous to combine them. Conversely, if their combination has some form of cost and a partial or total conflict exists, which we expect to be the generic case, then our results predict that there will be alternation and that it will follow the same logic identified here.

We conclude by noting that, in addition to the familiar cases of dogs and rodents mentioned in the introduction, other species can sense chemical cues both in the bulk and on surfaces, and may feature a similar phenomenology of alternation. In particular, a large body of experimental evidence has been collected for turbulent plume-tracking by aquatic organisms, as reviewed in [21]. Crustaceans sense chemical cues with their antennules floating in water and switch to sensing with their feet as they approach the target [22]. For example, lobsters were observed in dim light in a flume of dimensions 2.5m x 90cm x 20cm, as they left their shelter upon release of a turbulent plume of odor obtained from grounded mussel [23]. As the animals encountered the plume, they often displayed special behaviors, including raising up, sweeping their sensory legs on the bottom of the flume and increasing flicking of lateral antennules. Similar observations were made for blue crabs capturing live clams or tracking spouts releasing clam extract [24]. In these experiments blue crabs would occasionally lower their abdomen closer to the surface or extend their walking legs to raise above their normal height. Finally, pelagic marine mollusks *Nautilus pompilius* were observed to track the source of a turbulent plume by swimming at different heights, above and below the center of the plume. Interestingly, most animals sampled at higher heights beyond one meter from the source, and swam at lower heights when closer to the source [25]. These experiments indicate that animals may alternate between different heights, and that sampling at higher elevation may be particularly useful at larger distances, which is again in qualitative agreement with our results. The ensemble of these observations suggest that alternation between sensorimotor modalities is likely to be present in the behavior of aquatic organisms as well. We hope that results presented here will motivate more experiments, on dogs, rodents and aquatic organisms alike, with the goal of assessing quantitative aspects of the observed behaviors, testing our framework and advancing understanding of how sensorimotor modalities are integrated.

## ACKNOWLEDGMENTS

G.R was partially supported by the NSF-Simons Center for Mathematical & Statistical Analysis of Biology at Harvard (award number #1764269) and the Harvard Quantitative Biology Initiative. This work received support from: the European Research Council (ERC) under the European Union’s Horizon 2020 research and innovation programme (grant agreement No. 101002724 RIDING); the Air Force Office of Scientific Research under award number FA8655-20-1-7028; the National Institutes of Health (NIH) under award number R01DC018789. The authors are grateful to the OPAL infrastructure from Université Côte d’Azur and the Université Côte d’Azur’s Center for High-Performance Computing for providing resources and support. N.R. is thankful for the support of Instituto Nazionale di Fisica Nucleare (INFN) Scientific Initiative SFT. This research was initiated at the Kavli Institute for Theoretical Physics supported in part by NSF Grant No. PHY-1748958 and the Gordon and Betty Moore Foundation Grant No. 2919.02.

## MATERIALS AND METHODS

### Direct numerical simulations

The Navier-Stokes (M1) and the advection-diffusion equation for passive odor transport (M2) describe the spatiotemporal evolution of odor released in a fluid. We can solve these equations with direct numerical simulations (DNS) and obtain realistic odor fields to feed the POMDP algorithm:

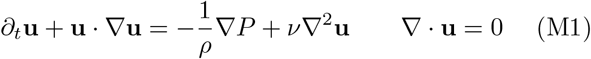

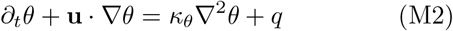

where **u** is the velocity field, *ρ* is the fluid density, *P* is pressure, *ν* is the fluid kinematic viscosity, *θ* is the odor concentration, *κ*_*θ*_ is its diffusivity and *q* an odor source.

We simulate a turbulent channel of length *L*, width *W* and height *H*, where fluid flows from left to right and hits a solid hemicylindrical obstacle of height 38 cm set on the ground, which produces turbulence. A horizontal parabolic velocity profile is set at the left boundary 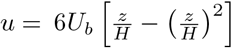, where *z* is the vertical coordinate and *U*_*b*_ is the mean horizontal speed. We impose the no-slip condition at the ground and on the obstacle and an outflow condition at the other boundaries (see [26] for more details).

When the turbulent flow is fully developed a concentrated odor source is added at 0.58 m from the ground, i.e. 20 cm above the center of the obstacle. The source is defined by a Gaussian profile with radius *σ* ∼ 5*η*, where *η* is the smallest scale of turbulent eddies (see Table S1). We set adsorbing boundary conditions at the inlet, on the ground and on the obstacle and zero gradient conditions on the sides and top.

The simulation was realized by customizing the open-source software Nek5000 [27] developed at Argonne National Laboratory, Illinois. The three dimensional volume of the channel is discretized in a finite number of elements and Nek5000 solves the Navier-Stokes and scalar transport equations within every element with a spectral element method. To accurately describe all relevant scales of turbulence from the dissipative scale to the length of the domain, the solution is expanded in 8th grade polynomials in each of 160 000 elements, thus effectively discretizing space in 81 920 000 grid points. Table S1 summarizes the parameters that characterize the flow. Each DNS runs for 300 000 time steps where 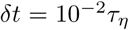 following a strict Courant criterium with *U*Δ*t/*Δ*x* < 0.4 to ensure convergence of both the velocity and scalar fields. Snapshots of velocity and odor fields are saved at constant frequency *ω* = 1*/τ*_*η*_. Fully parallelized simulations require 2 weeks of computational time using 320 cpus, see ref. [26] for further details.

### The POMDP framework

We briefly introduce Partially Observable Markov Decision Processes (POMDPs) before describing the specific algorithms used in our simulations. We refer to [28] for a detailed review on POMDPs. POMDPs are a generalization of Markov Decision Processes (MDP) analogous to the relationship between Hidden Markov models and Markov models [19, 20]. In an MDP, we define a state space, an action space and a reward function. The dynamics of the state space is Markovian and is defined entirely by the transition matrix, *T* (*s*′|*s, a*), which gives the probability of transitioning to state *s*′ given the current state *s* and the action taken, *a*. After each transition, the agent receives a reward, which has expectation *r*(*s, a, s*′). Given the transition matrix and the reward function, the goal is typically to find the unique optimal policy, Π*(*s*), which maximizes the discounted sum of future rewards, ⟨*r*_0_ + *γr*_1_ + *γ*^2^*r*_2_ + … ⟩, where *γ* is the discount factor and *r*_*t*_ is the expected reward *t* steps after the initial state. Often, but not always, this involves solving for the value function, *V* (*s*), which is the expected discounted sum of rewards from state *s*, conditional on policy Π*(*s*). The value function satisfies the central dynamic programming equation known as the Bellman equation [29]:

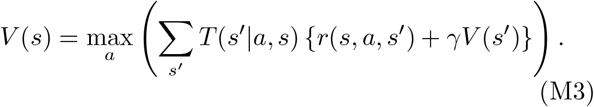

While MDPs deal with fully observable states, POMDPs have one additional feature which makes it appropriate for our setting. Instead of observing the current state, the agent only receives certain observations, *o*, from which the true latent state has to be dynamically inferred. The agent is assumed to have a model of the environment, *P* (*o*|*s, a*). In our setting, this likelihood function encodes the statistics of detections and non-detections at various locations down-wind of an odor source (Figure S3). A POMDP therefore maps a sequence of recent observations and actions *o*_−1_, *a*_−1_, *o*_−2_, *a*_−2_, … to a strategic action. While the dimensionality increases rapidly with the length of the observation history, the entire history is encoded by the current posterior distribution over states, ***b***, also known as the *belief vector*. The problem of solving for the optimal action is recast as the problem of solving for the policy Π*(***b***). The Bellman equation on states for MDPs translates into a Bellman equation on belief vectors for POMDPs:

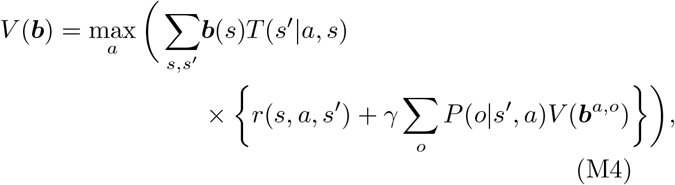

where ***b***^*a,o*^ is the posterior belief state given the agent takes action *a* and observes *o*. Using Bayes’ rule, ***b***^*a,o*^ is given by

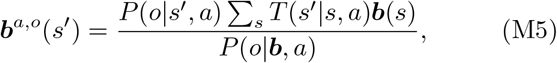

where the normalizing factor is

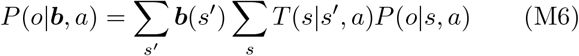

Intuitively, the Bayes’ rule takes into account the new information gained from the most recent observation and the information lost due to the state space dynamics, which are in turn influenced by the action.

### Algorithms to solve POMDPs

We use POMDP-solvers which approximate the value function, *V* (***b***), for all ***b***. If the value function is known, the optimal policy is simply to choose the action that yields the highest future expected return given the current belief vector :

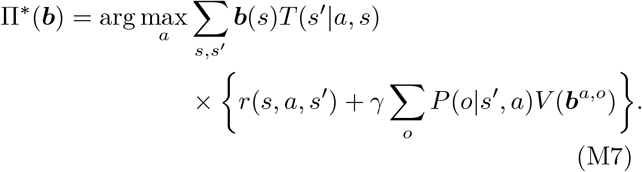

Computing the value function *V* (***b***) exactly for all belief vectors for tasks containing more than a handful of states is infeasible. Existing methods exploit a specific representation of the value function, which leads to the approximation discussed by [28]. We recapitulate here the main results and refer to [28] for more details. In particular, it can be shown that the value function can be approximated arbitrarily well by a finite set *H* of hyperplanes [30], each of which is parameterized by ***α***(*s*):

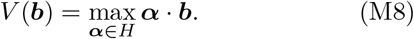

An initial set *H* is expanded using the Bellman equation (M4). Using vector notation, we can write

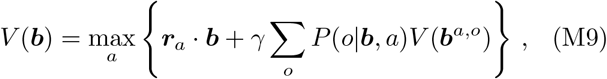

where ***r***_*a*_(*s*) = Σ_*s*_^*′*^ *T* (*s*′|*a, s*)*r*(*s, a, s*′). Let ***α***^*a,o*^(***s***) be defined as

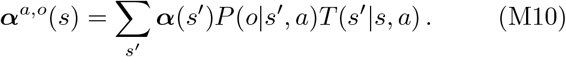

and using the belief update (M5) it follows that

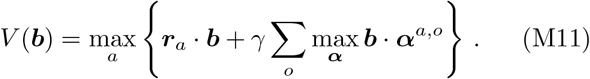

Given the previous set *H*, we can add a new ***α*** vector to it corresponding to belief vector ***b*** called the “backup” operation:

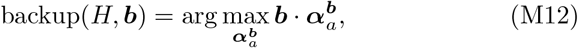

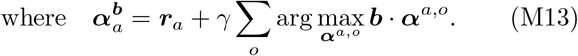

In other words, given a previous set *H* and new belief vector, one can use the Bellman equation to update *H* and obtain a better approximation to the value function. The key computational advantage of using the above backup operation is that the ***α***^*a,o*^’s can be pre-computed for the current *H* and re-used when backing up.

The question then is: how do we efficiently collect new belief vectors to update *H* and prune vectors from *H* that are no longer necessary? Algorithms differ at these two stages. We use Perseus [31], which simulates random exploration of the agent. Specifically, at each step in a “training” episode, we start from an initial prior, pick actions (uniform) randomly and then sample observations from *P* (*o*|***b***, *a*). The new belief vector obtained using Bayes’ rule is then used to backup *H*. Finally, after adding a new set of ***α*** vectors into *H*, it is efficient to prune the existing ones that are guaranteed to not be used. We prune the ***α*** vectors whose every component is smaller than those of another vector (see [28] for other heuristic pruning methods).

Three parameters can be tuned: the discount factor *γ*, the number of belief points sampled per each episode of random exploration and the total number of training episodes. The discount rate sets the planning horizon, which is set to be of the same order as the typical number of steps to get to the target. Increasing the latter two parameters improves the strategy at the expense of increased training time. We solve the POMDP for various values of these two parameters and show that the performance saturates at a parameter range within computational feasibility.

### POMDPs for learning sniff-and-search strategies

To implement POMDPs that learn to navigate odor plumes by employing multiple modes of search, we consider a simple state space consisting of a 2d grid with dimensions 10 × 2 discretized with 30 points per unit length so that the state space has size 10 × 2 × 30^2^ = 18, 000. The agent can take 6 possible actions corresponding to movement in either of the four directions or staying at the same spot while sniffing ground odor cues. The sixth action corresponds to staying at the same spot and sniffing airborne odor cues. After every action, the agent can make one of three observations – no detection, an odor detection or finding the odor source. Odor detections are binarized, i.e., the odor is detected if the concentration is above a certain threshold. The intermittent nature of turbulent fluctuations imply that there is little additional information in the graded concentration beyond the information contained in the detection rate [13, 32]. We assume a Poisson rate of detection with the rate map at the ground level and at the nose level when sniffing in the air obtained by measuring the fraction of time the odor concentration is above 0.14% with respect to the maximum concentration at source in the flow simulations. As described above, we use the Perseus algorithm [31], which performs random exploration starting from a given prior belief of where the source is located. We use a uniform prior of dimensions 28.6 *m* × 3.4 *m*. After training, the POMDP algorithm yields a set *H* which encodes an approximation to the value function mapping belief vectors to expected discounted rewards for each of the possible actions. The decision at each step is then obtained from (M7).

### Parameters for POMDP used in main text

The main figures represent results using: discount factor *γ* = 0.99, number of training episodes *i* = 320, number of belief points sampled per training episode *i*′ = 100, likelihood in the air and at the ground is defined as shown in Figure S3, in Figure 6 likelihood in the air is defined as a rectangle with dimensions *x*_thr_ × *y*_thr_. Results in Figure 4 are tested and averaged over three different starting positions (x = 8; y = 0, 0.3, −0.5), 8 different seeds, 50 different realizations for the same seed (trajectories differ for the history of detections according to the Poissonian model).

The effect of varying *γ* are represented in Figure S2 (all other parameters are kept constant); the effect of varying the number of training episodes is represented in Figure S1 (averaged over 3 different locations and 3 seeds).

Training requires up to 2 days in time on 1 processor, while testing a single realization takes ∼ 10 hours.

### Derivation of (2)

We consider a scenario where a target is located at one of *N* possible patches, *n* = 1, 2, …, *N* with probabilities ***p***_0_ = (*p*_1_, *p*_2_, …, *p*_*N*_) (Σ_*n*_ *p*_*n*_ = 1). Note that *p*_*n*_ = 1*/N* for all *n* for the prior considered in the main text. The agent starts at *n* = 1 and moves sequentially from *n* = 1 to *n* = *N* while spending time *t*_*n*_ sampling in each patch. Moving from a patch to the next one takes time *τ* ≡ *x*_thr_*/v*. At *n* = *N*, the agent samples indefinitely, *t*_*N*_ = ∞. The agent receives reward of one when the target is found in a patch, which is discounted at rate *λ*. The value *V*_1_ ≡ ⟨*e*^−*λT*^ ⟩_*T*_, where *T* is the search time, is the expected discounted reward optimized w.r.t *t*_*n*_’s. We derive two sets of recursive equations (with and without Bayesian updates) to calculate *V*_1_. We show that both formulations lead to the same optimal casting times, however, the set of equations without Bayesian updates are much simpler to compute.

Suppose the cumulative probability of finding the target in time *t conditional* on the target being in that patch is *d*(*t*). Note that *c*(*t*) ≡ 1 − *d*(*t*) is used in the main text. Denote 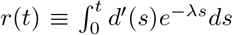. This is the expected discounted reward if the agent searches for time *t* in a patch that contains the target.

Say *N* = 3. Since *V*_1_ is the expected discounted reward optimized over the casting times *t*_1_, *t*_2_, we have

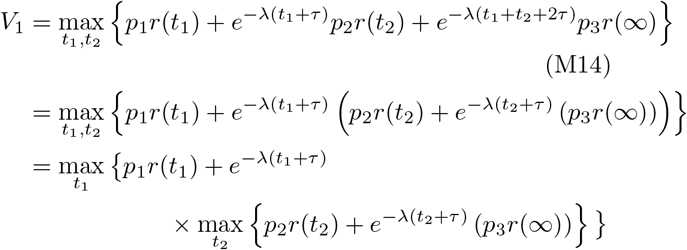

The last equation above motivates a recursive equation for general *N* :

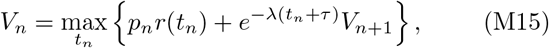

with boundary condition, *V*_*N*_ = *p*_*N*_ *r*(∞). Optimizing over *t*_*n*_, we obtain the marginal value condition

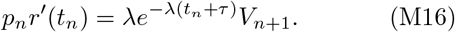

If the rate of detection during casting is a constant *κ*, we have 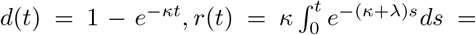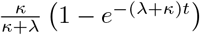 and *r*′(*t*) = *κe*^−(*κ*+*λ*)*t*^. Plugging this expression into (M16), we get

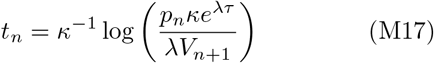

Now, let’s calculate *V*_1_ using Bayesian updates and show that the optimal times exactly correspond to what we have in the previous equation. Denote 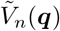 as the value at patch *n* for an arbitrary probability vector ***q*** = (*q*_1_, *q*_2_, …, *q*_*N*_). We now show that 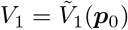, where ***p***_0_ = (*p*_1_, *p*_2_, …, *p*_*N*_) is the prior. We have

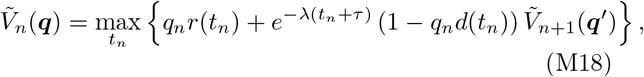

where ***q***′ is the posterior conditional on no detection. The two terms on the r.h.s correspond to the case when the agent finds the target in the patch before *t*_*n*_ (with probability *q*_*n*_*d*(*t*_*n*_)) and does not find it (with probability 1 − *q*_*n*_*d*(*t*_*n*_)) respectively. Given the observation that the target is not found in patch *n*, the posterior probabilities, ***q***′, are obtained using Bayes’ rule:

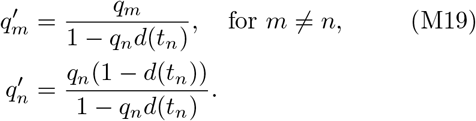

We show that 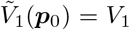 for *N* = 3. The general case of starting from any patch, prior and number of patches (*N*) follows. Expanding (M18) starting from *n* = 1,

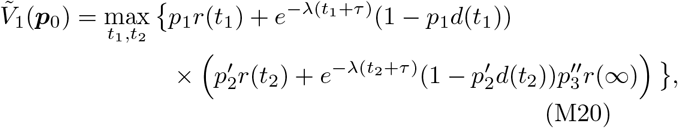

where 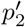 is obtained from the first Bayesian update and 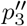 is obtained after the second Bayesian update. Using (M19), we have 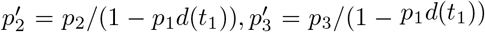 and 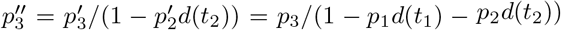.

Since 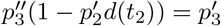, simplifying (M20), we get

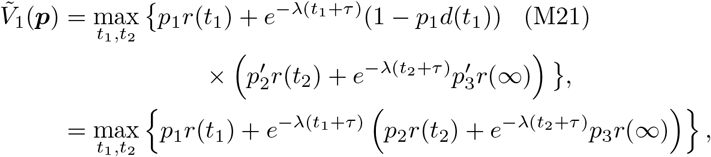

where 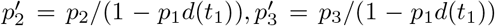 are used in the second step. This equation exactly corresponds to (M14). The upshot is that the normalization factors from the Bayesian updates go through the parenthesis and cancel out. However, optimizing for *t*_*n*_ directly using (M18) is difficult due to the dependence of ***q***′ on *t*_*n*_.

---

**TABLE S1.** Parameters of the simulation. Length *L*, width *W*, height *H* of the computational domain; horizontal speed along the centerline *U* ; mean horizontal speed *U*_*b*_ = ⟨*u*⟩; Kolmogorov length scale *η* = (*ν*^3^*/ϵ*)^1*/*4^ where *ν* is the kinematic viscosity and *ϵ* is the energy dissipation rate; mean size of gridcell Δ*x*; Kolmogorov timescale *τ*_*η*_ = *η*^2^*/ν*; energy dissipation rate 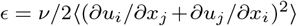; Taylor microscale 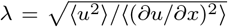; wall lengthscale *y*^*+*^ = *ν/u*_*τ*_ where the friction velocity is 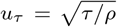 and the wall stress is *τ* = *ρνdu/dz* |_*z*=0_; Reynolds number *Re* = *U*(*H/*2)*/ν* based on the centerline speed *U* and half height; Reynolds number *Re*_*λ*_ = *Uλ/ν* based on the centerline speed and the Taylor microscale *λ*; magnitude of velocity fluctuations *u*′ relative to the centerline speed; large eddy turnover time *T* = *H/*2*u*′. First row reports results in non dimensional units; second row corresponds to dimensional parameters in air, assuming the mean speed is 25 cm/s.

| $L$ | $W$ | $H$ | $U$ | $U_b$ | $\eta$ | $\Delta x$ | $\tau_\eta$ | $\epsilon$ | $\lambda$ | $y^+$ | Re | $Re_\lambda$ | $u'/U$ | $T$ |
| --- | --- | --- | --- | --- | --- | --- | --- | --- | --- | --- | --- | --- | --- | --- |
| 40 | 8 | 4 | 32 | 25 | 0.006 | 0.025 | 0.01 | 39 | 0.17 | 0.004 | 16000 | 1370 | 10% | $64\tau_\eta$ |
| 15 m | 3 m | 1.5 m | 0.33 m/s | 0.25 m/s | 0.23 cm | 1 cm | 0.36 s | $1.2e-4 \text{ m}^2/\text{s}^3$ | 6 cm | 0.14 cm | | | | |

**FIG. S1.**
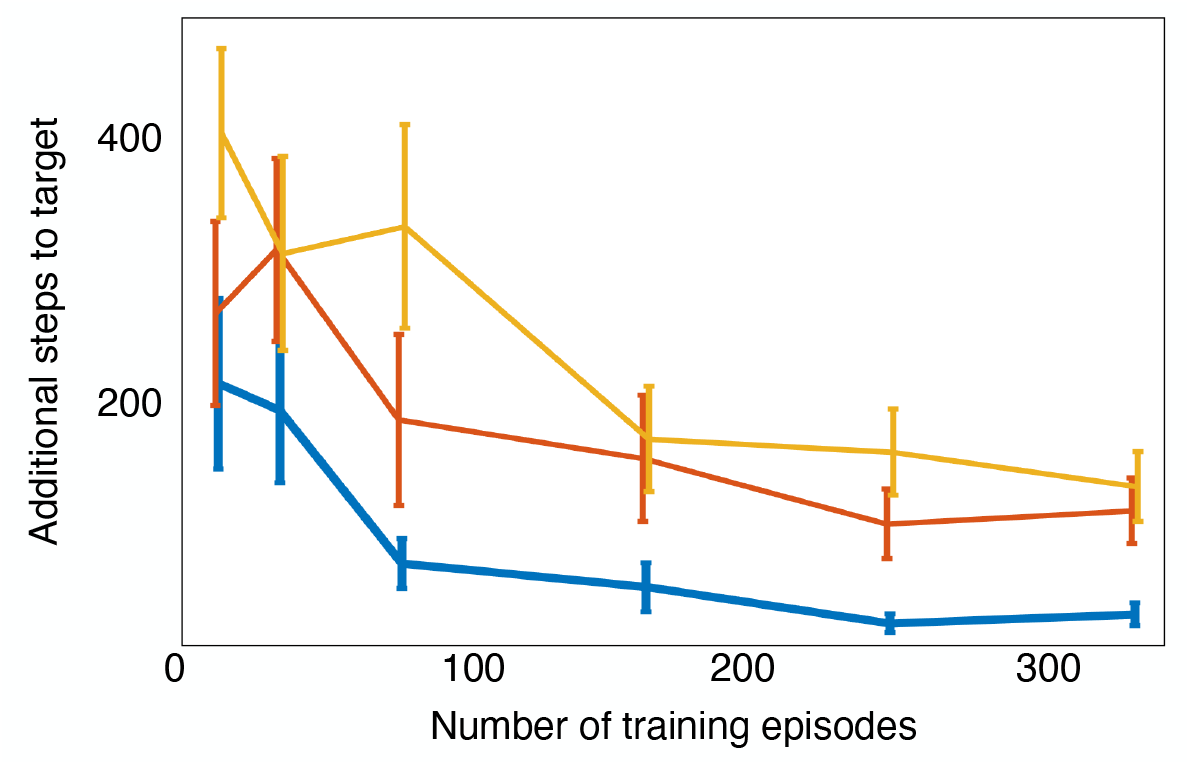
Performance of POMDP improves with training episodes and saturates after a certain number of iterations. Training converges for different values of the discount factor *γ*, yellow is *γ* = 0.90, red is *γ* = 0.95 and blue is *γ* = 0.99. The gain in performance with increasing *γ* is indicative of the development of a long-term strategy. We use *γ* = 0.99 and number of training episodes *i* = 320 throughout the Results section.

**FIG. S2.**
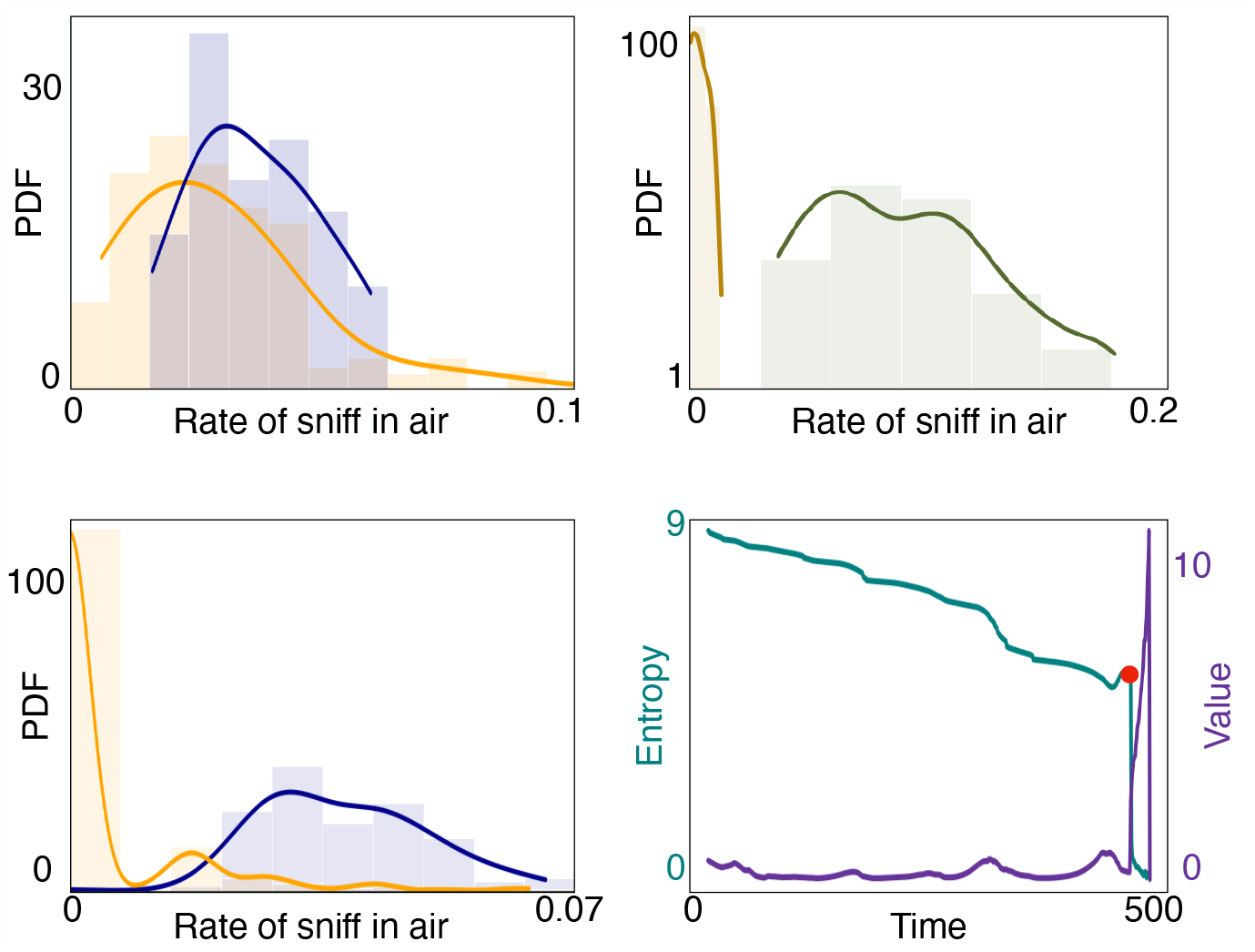
Here we show the results analogous to Figure 4 with a smaller discount factor *γ* = 0.95. Alternation between olfactory modalities is preserved, as well as surging and casting.

**FIG. S3.**
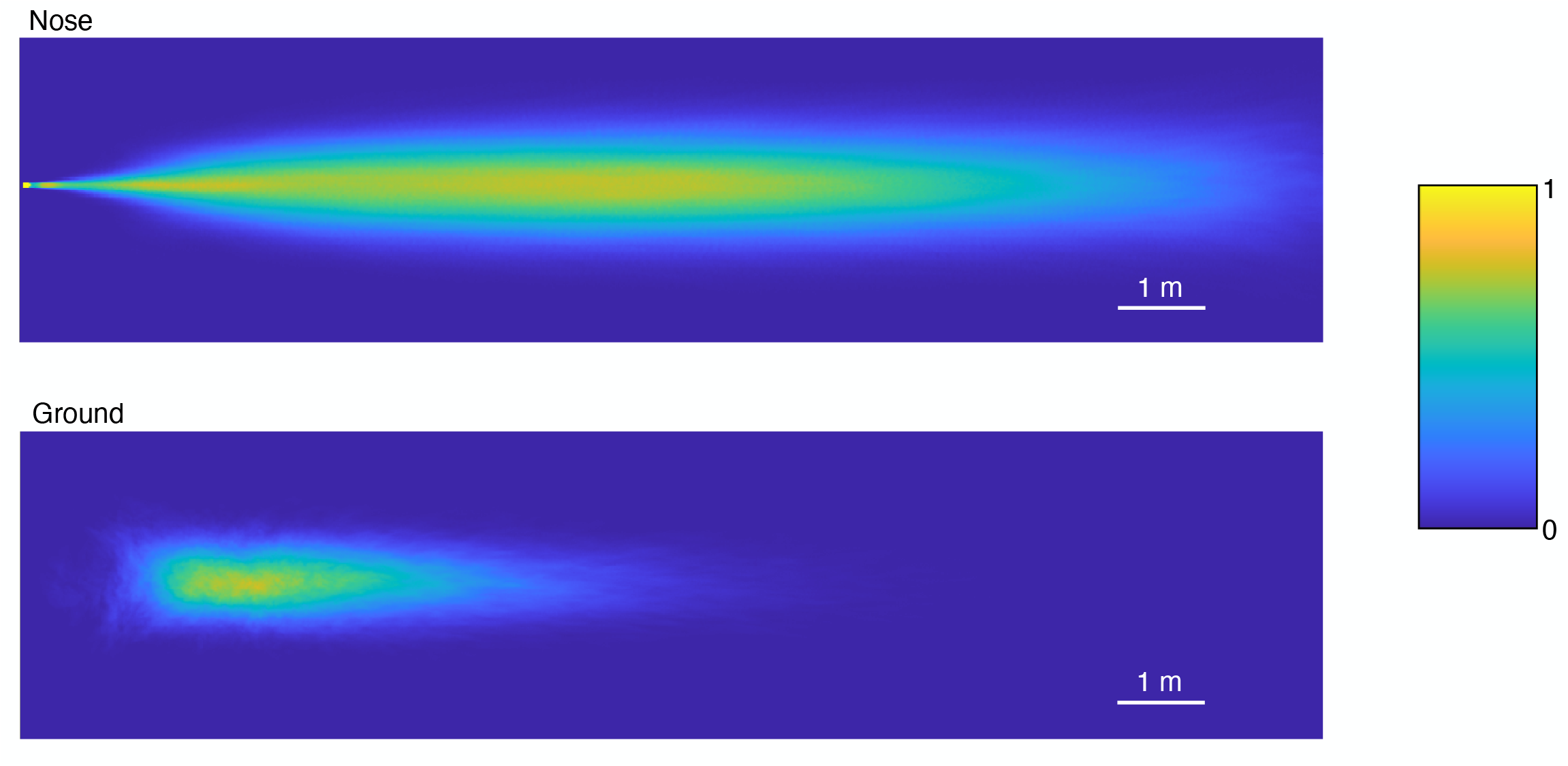
The probability per unit step of detecting an odor signal in the air and at the ground obtained from direct numerical simulations of odor transport. These detection rate maps constitute the observation likelihood models used to train the POMDP. Note that the arena defined in the POMDP is larger than the rate maps shown here (see Figure 3 for instance). The detection rate is set zero beyond the bounds of the above rectangles.

